# Evaluating FAIR Maturity Through a Scalable, Automated, Community-Governed Framework

**DOI:** 10.1101/649202

**Authors:** Mark D Wilkinson, Michel Dumontier, Susanna-Assunta Sansone, Luiz Olavo Bonino da Silva Santos, Mario Prieto, Dominique Batista, Peter McQuilton, Tobias Kuhn, Philippe Rocca-Serra, Mercè Crosas, Erik Schultes

**Author notes:** Corresponding authors: Mark Wilkinson, Susanna-Assunta Sansone, Erik Schultes.

## Abstract

Transparent evaluations of FAIRness are increasingly required by a wide range of stakeholders, from scientists to publishers, funding agencies and policy makers. We propose a scalable, automatable framework to evaluate digital resources that encompasses measurable indicators, open source tools, and participation guidelines, which come together to accommodate domain relevant community-defined FAIR assessments. The components of the framework are: (1) Maturity Indicators - community-authored specifications that delimit a specific automatically-measurable FAIR behavior; (2) Compliance Tests - small Web apps that test digital resources against individual Maturity Indicators; and (3) the Evaluator, a Web application that registers, assembles, and applies community-relevant sets of Compliance Tests against a digital resource, and provides a detailed report about what a machine “sees” when it visits that resource. We discuss the technical and social considerations of FAIR assessments, and how this translates to our community-driven infrastructure. We then illustrate how the output of the Evaluator tool can serve as a roadmap to assist data stewards to incrementally and realistically improve the FAIRness of their resources.

## Introduction

The FAIR Data Principles^1^ were designed to ensure that all digital resources can be Findable, Accessible, Interoperable, and Reusable by machines. The Principles act as a guide to the kinds of behaviours that researchers and data stewards should increasingly expect from digital resources^2^, and in turn, impose compliance-requirements on researchers if they wish to publish scholarly outputs FAIRly. In recent years, funding agencies, journals, community alliances, learned societies and research organizations both public and private have begun to endorse the creation and publication of FAIR digital resources.

The FAIR Principles are aspirational, in that they do not strictly define how to achieve a state of “FAIRness”, rather they describe a continuum of features, attributes, and behaviors that move a digital resource closer to that goal. Despite their rapid community uptake, the question of how the FAIR Principles should be implemented has been prone to diverse interpretation. Some resource providers claim to be “already FAIR” or “to enable FAIRness” - statements that currently cannot be objectively evaluated. These manifold interpretations of the FAIR Principles are counterproductive, posing a risk of fragmentation and confusion in a manner antithetical to their intended purpose (or worse, being ignored entirely due to a lack of formal clarity). This state of affairs is somewhat ironic in that, because FAIR speaks to machine-actionable operations, FAIR digital objects should therefore be amenable to unambiguous and indeed, completely automated forms of validation and evaluation.

In the following sections we describe the development path of our proposed scalable, automatable FAIRness evaluation framework, and how each step has been informed by community input and feedback. We also highlight the unique features of our approach, particularly those that distinguish this work from manually-executed, questionnaire-based evaluations.

### Our community-driven, phased approach

Early discussions with stakeholder communities highlighted some important behaviors or features that should be reflected in, or result from, an initiative aimed at FAIR evaluation. In particular:

1. Some FAIR behaviors may be considered ‘universal’ but these may be complemented by additional resource-specific behaviors that reflect the expectations of particular communities.
2. The behavioural indicators themselves, their tests, and any results stemming from their application, must be FAIR.
3. Use of open standards for all components should foster a vibrant ecosystem of FAIRness assessment tools.
4. Various approaches to FAIRness assessment should be enabled (e.g. self-assessment, task forces, crowd-sourcing, and automated), however, the ability to scale FAIRness assessments to billions if not trillions of diverse digital objects is critical.
5. FAIRness assessments should be kept up to date, and all assessments should be versioned, have a time stamp and be publicly accessible.
6. Consideration should be given to simple visualizations of outputs, as these will be a powerful modality to inform users and guide the work of producers of digital assets.
7. The assessment process should be designed and disseminated to positively incentivise the digital resource, which should benefit from this and use it as an opportunity for improvement.
8. Maximizing an evaluation score should not be a goal in and of itself, nor is there any intrinsic value in evaluations; rather they should assist in meaningful cost/benefit analysis for FAIRification efforts.

With these guiding principles in-mind, an open framework^3^ for the creation and publication of Metrics for FAIRness was designed, together with an initial set of 14 exemplar FAIR Metrics. These described specific expectations of FAIR resources, and possible approaches to checking compliance. The 14 Metrics were used in an initial study of 11 key data resources. Referred to as “Generation 1” (and hereafter “Gen1”), the Metrics were formulated into a series of questions, provided as a questionnaire to the resource owners or users familiar with the resource. The responses to these questionnaires were then manually evaluated as a means to measure the utility and clarity of the Metrics themselves (rather than evaluating the resource to which they were applied). The outcomes from this study were returned to the participants for their feedback, and compiled into a self-archived report^4^. In the next sections we summarize the lessons learned from this first phase of the work, and how we have benefitted from this community feedback, both positive and negative, to evolve the framework.

Admittedly, there was some resistance from data and resource providers who were concerned about how the results of these FAIRness assessments would affect them, especially when the tests were carried out by third parties and without their knowledge or participation. We have been very sensitive to those concerns, and as a result we worked with willing individuals and groups to revise our approach. The Gen1 Metrics fell-short of their objectives in a variety of ways: they were designed with insufficient awareness of practices being used within the data publishing community, encouraged overly simple responses, defied simple validation, and were easily misunderstood^4^; moreover, the word ‘metrics’ itself seemed off-putting.

With the desire to pursue a more objective assessment framework, we conducted a deeper exploration of the data provision “ecosystem” to better understand what the data provider community was doing *vis a vis* what *they* believed were FAIR practices. This led to a deeper understanding of the global state of FAIRness, and the surprisingly wide range of in-use practices we found with respect to (meta)data provision. Based on this, a second round of authoring was undertaken. To address the Gen1 flaws, we replaced the term “Metrics” with “Maturity Indicators” (MIs), where MIs describe facets of FAIRness that can be objectively evaluated by a machine and thus used to establish a “contract of expectations and capabilities” between a data resource and an automated agent. For example, one of the MIs for FAIR Principle F1 (Findability) evaluates if a repository implements structured metadata to enhance its discoverability on the Web; a second MI evaluates if this metadata follows a community-defined model. Thus, through these two MIs, an agent that requires metadata following the rules of Linked Data could pre-determine the capability of that specific repository to support that requirement, before attempting to retrieve and utilize it.

Guided by the community feedback, we have evolved the framework from a questionnaire-based approach to an automatable and we believe, scalable framework, which utilizes a computational agent to autonomously discover, access, and interpret the content of a data resource. Provided with only the Globally Unique IDentifier (GUID) of the metadata for the digital resource to be assessed, the FAIR Evaluator, executes a set of MI tests and generates a detailed report regarding the features of FAIRness it discovered while exploring that resource, and how it made those determinations. To ensure that the MIs are driven by the needs of communities and the specifics of the digital objects to be assessed, we established a well-documented and open authoring framework (https://github.com/FAIRMetrics/Metrics), such that all stakeholders may participate in the creation of domain-relevant community-specific MIs. As exemplars, we authored a starter set of 15 “bootstrap” MIs, and their associated automated tests (designated as “Generation 2” or “Gen2”) that may act as starting-points for others to follow or extend.

In the following sections we detail the design and the components of our scalable, automatable framework for the objective and quantitative evaluation of the level of FAIRness of digital objects. We also illustrate how the enhanced, open infrastructure puts FAIR assessment in the hands of the community, while also serving as a roadmap to incremental improvements in the FAIRness of a resource.

## Results

The design considerations for our FAIRness evaluation framework were:

1. Anyone should be able to evaluate any digital object.
2. Anyone, as an individual or preferably as a community, can create and publish a MI or a collection of MIs.
3. Evaluations should use domain-relevant, community-defined Compliance Tests.
4. All components of the evaluation framework should themselves be FAIR.
5. The evaluation should be as objective as possible; the evaluation code should avoid accommodating non-standards-compliant behaviors wherever possible.
6. The evaluation should provide the user not just with the result, but also with clear guidance for improving the level of FAIRness.
7. The evaluation framework should be modular and extendable.
8. The framework should automatically update to accommodate new standards or technologies whenever possible, to minimize maintenance and sustain utility over time.

### Starter pack

A set of 15 MIs were defined, using the template described in the Methods section (Box 1). These cover most of the FAIR Principles and sub-principles, with the exception of R1.2 and R1.3. MIs are identified by a GUID which resolves to the nanopublication^5^ representing the machine-readable record of the MI, such that the MIs themselves are FAIR digital objects.

#### Box 1. Bootstrap FAIR MIs.

“Loose” and “strict” is an indication of the rigor with which the MI says a resource should be tested. For example, “Uses FAIR vocabularies (loose)” indicates that it will look for Linked Data in the resource, and then test if the predicates resolve (to anything) by HTTP protocols. “Uses FAIR vocabularies (strict)” further requires that these predicates resolve to Linked Data, using content-negotiation for Linked Data formats.

1. Identifier uniqueness (F1)
2. Identifier persistence (F1)
3. Structured metadata (F2)
4. Grounded metadata (F2)
5. Use of GUIDs in metadata (F3)
6. Metadata indexed in a searchable resource (F4)
7. Open protocol for (meta)data retrieval (A1.1)
8. Protocol supports authentication/authorization (A1.2)
9. Metadata persistence (A2)
10. Use a knowledge representation language (loose) (I1)
11. Use a knowledge representation language (strict) (I1)
12. Uses FAIR vocabularies (loose) (I2)
13. Uses FAIR vocabularies (strict) (I2)
14. Qualified outward links (I3)
15. Metadata contains link to license (R1.1)

For each of the bootstrap MIs, at least one Compliance Test was written to evaluate a (meta)data resource’s adherence to that MI. These are published as standalone Ruby Web applications, with interfaces defined using FAIR smartAPI^6,7^-based metadata, as described in the Methods section. The descriptive names of these tests are listed and numbered in Box 2, together with the GUID of the MI to which they are associated.

#### Box 2: Bootstrap Compliance Tests

1. Unique identifier (https://w3id.org/fair/maturity_indicator/terms/Gen2/Gen2_MI_F1A)
2. Data identifier explicitly in metadata (https://w3id.org/fair/maturity_indicator/terms/Gen2/Gen2_MI_F3)
3. Metadata identifier explicitly in metadata (https://w3id.org/fair/maturity_indicator/terms/Gen2/Gen2_MI_F3)
4. Identifier persistence (https://w3id.org/fair/maturity_indicator/terms/Gen2/Gen2_MI_F1B)
5. Data identifier persistence (https://w3id.org/fair/maturity_indicator/terms/Gen2/Gen2_MI_F1B)
6. Structured metadata (https://w3id.org/fair/maturity_indicator/terms/Gen2/Gen2_MI_F2A)
7. Grounded metadata (https://w3id.org/fair/maturity_indicator/terms/Gen2/Gen2_MI_F2B)
8. Searchable in major search engine (https://w3id.org/fair/maturity_indicator/terms/Gen2/Gen2_MI_F4)
9. Uses open free protocol for data retrieval (https://w3id.org/fair/maturity_indicator/terms/Gen2/Gen2_MI_A1.1)
10. Uses open free protocol for metadata retrieval (https://w3id.org/fair/maturity_indicator/terms/Gen2/Gen2_MI_A1.1)
11. Data authentication and authorization (https://w3id.org/fair/maturity_indicator/terms/Gen2/Gen2_MI_A1.2)
12. Metadata authentication and authorization (https://w3id.org/fair/maturity_indicator/terms/Gen2/Gen2_MI_A1.2)
13. Metadata persistence (https://w3id.org/fair/maturity_indicator/terms/Gen2/Gen2_MI_FA2)
14. Metadata knowledge representation language (weak) (https://w3id.org/fair/maturity_indicator/terms/Gen2/Gen2_MI_I1A)
15. Metadata knowledge representation language (strong) (https://w3id.org/fair/maturity_indicator/terms/Gen2/Gen2_MI_I1B)
16. Data knowledge representation language (weak) (https://w3id.org/fair/maturity_indicator/terms/Gen2/Gen2_MI_I1A)
17. Data knowledge representation language (strong) (https://w3id.org/fair/maturity_indicator/terms/Gen2/Gen2_MI_I1B)
18. Metadata uses FAIR vocabularies (weak) (https://w3id.org/fair/maturity_indicator/terms/Gen2/Gen2_MI_I2A)
19. Metadata uses FAIR vocabularies (strong) (https://w3id.org/fair/maturity_indicator/terms/Gen2/Gen2_MI_I2B)
20. Metadata contains qualified outward references) (https://w3id.org/fair/maturity_indicator/terms/Gen2/Gen2_MI_I3A)
21. Metadata includes license (strong) (https://w3id.org/fair/maturity_indicator/terms/Gen2/Gen2_MI_R1.1)
22. Metadata includes license (weak) (https://w3id.org/fair/maturity_indicator/terms/Gen2/Gen2_MI_R1.1)

There is not a one-to-one correspondence between the Compliance Tests and the MIs. For example, there are two tests for the MI “Gen2_MI_F3” (Use of GUIDs in metadata): the first, “Metadata identifier explicitly In metadata”, checks the metadata record to ensure it contains its own GUID, the second - “Data identifier explicitly In metadata” - assesses the presence of the data’s GUID within the metadata record.

Compliance Tests may be grouped into customizable Collections, which allow communities and stakeholders to create their own domain-relevant set(s) that emphasize different MIs according to their own views and needs. For example, a funding agency may want to ensure that, first and foremost, the license metadata is specified to clarify the terms-of-use for a dataset; conversely, for a data integration project the data-level interoperability may be prioritized. Collections are described with, and searchable via, metadata tags provided by the community that has created them. Collections of MIs can also be shared by several communities; this will foster, but not enforce, reuse of Collections, to promote consistency.

### Exemplar evaluations

Here we walk-through the execution of an evaluation by first creating a Collection of Compliance Tests, and then applying it. The targeted resource is the front-end Web application of the FAIR Evaluator itself, therefore a digital object of type “software”, rather than “data”. For the FAIRness assessment of a software application, the relevant Compliance Tests are only those that refer to features of the metadata, rather than the data (which in this case is the source code).

The first step to create a new Collection in the Evaluator registry is to select the relevant Compliance Tests from the drop-down list. For FAIR assessment of a software record, we will exclude tests 16 and 17, which are related to data knowledge representation languages and therefore not relevant to this case. We then describe our Collection adding relevant metadata tags, including a title, e.g. “Maturity Indicator Tests applicable to Software Applications”, a human-readable description, and contact information via an ORCiD. Upon submission to the Evaluator registry, our new Collection receives a persistent GUID and it is now ready for use in an evaluation.

To begin the evaluation, we select this Collection and enter the GUID of our software (https://fairsharing.github.io/FAIR-Evaluator-FrontEnd/). Depending on the number of tests in the Collection, the process may take several minutes, because the tests exhaustively try all paths to find and extract the necessary metadata via HTTP calls, potentially with high latency. A screenshot of the results of this evaluation is shown in Fig. 1, where we are informed that 10 out of 20 tests were successful.

**Figure 1:**
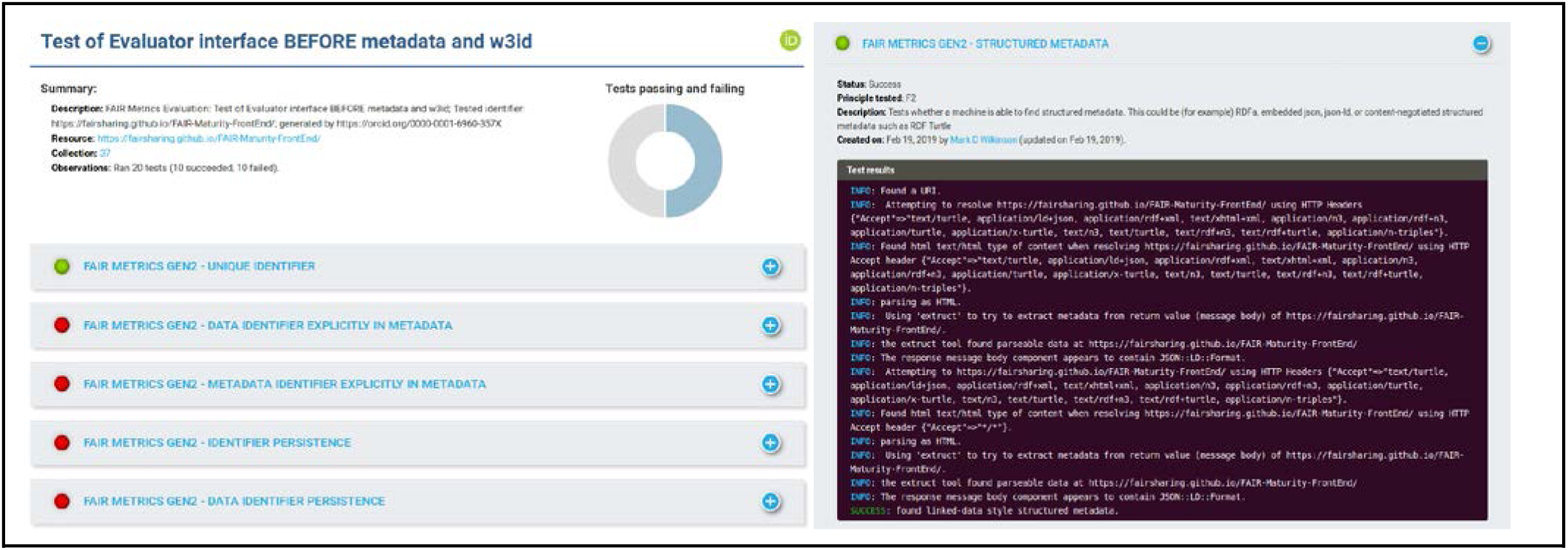
The output of assessment of the FAIR Evaluator’s interface. A summary and graph appear at the top of the report, followed by the results of individual tests as panels, with simple green (success) red (failure) indicators on each panel. Opening one of the result panels reveals an extensive, color-coded log of every activity undertaken by the Compliance Test, including the Metadata Harvester, and whether these ended in success or failure (and why). This allows the user to quickly scan the output for details, which will also serve as a guide to understand what it takes to improve the FAIRness of their resource.

### Turning evaluation into guidance

The information in this assessment is designed to guide users to improve the FAIRness of their digital objects. In our case, the Evaluator has reported failure on 10 Compliance Tests. Now, we explain how we used the output of these tests to improve the FAIRness of our software.

The result of “Metadata identifier persistence” (Fig. 2) informs us that, in order to pass this test, the GUID of the FAIR Evaluator page needs to be improved by adopting a community-recognised identifier schema, such as one of those listed by FAIRsharing (https://fairsharing.org/standards/identifierschema). Therefore we registered a new namespace for the FAIR Evaluator (AmIFAIR) with the w3id redirection system and obtained a permanent URL: https://w3id.org/AmIFAIR. This can now be referred-to in a metadata record for this software.

**Figure 2.**
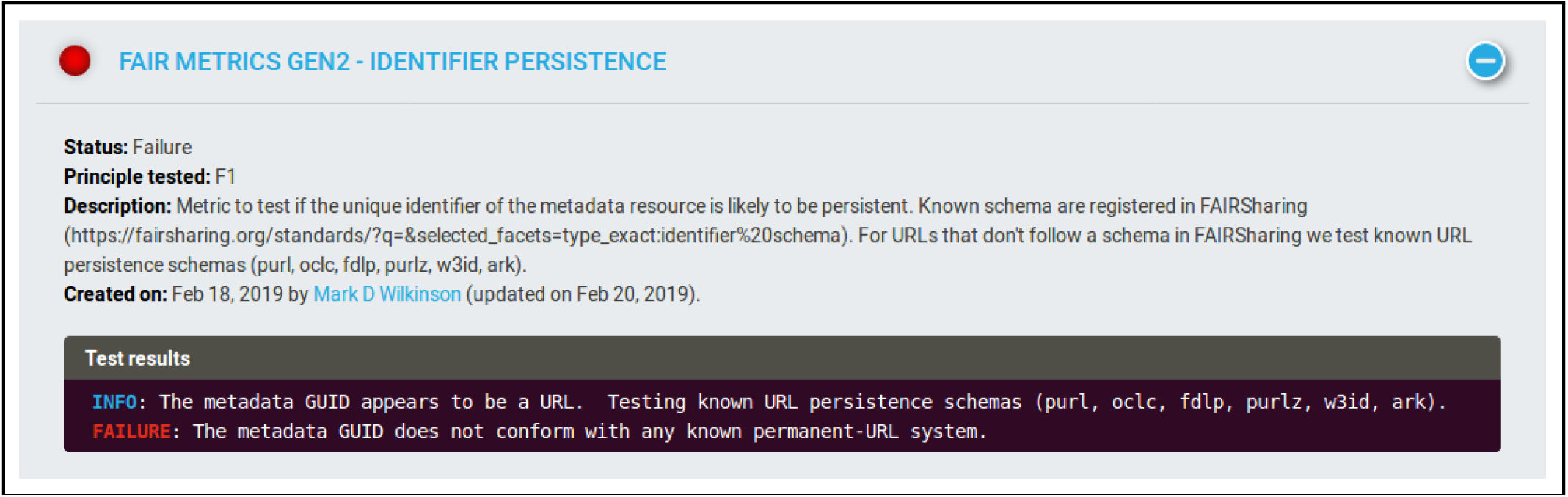
The output of the “Metadata identifier persistence” test. The information in the output indicates precisely what was observed (that the GUID presented to the test is a URL) and what is tested (compliance with known permanent-url schemas).

The information output from the “Metadata uses FAIR vocabularies (weak)” test (Fig. 3) informs us that the test received HTML when requesting Linked Data. The HTML that was provided in response to this HTTP request was therefore scanned by Extruct (https://github.com/scrapinghub/extruct) - a tool capable of extracting various kinds of embedded metadata from HTML. In this case, the report tells us that examination of the Extruct tool output identified some JSON-LD (this is metadata automatically generated by the Extruct tool itself in its output report, and has only locally-scoped URIs that are therefore ignored by the test); however, no author-provided metadata was found in the HTML of the page.

**Figure 3.**
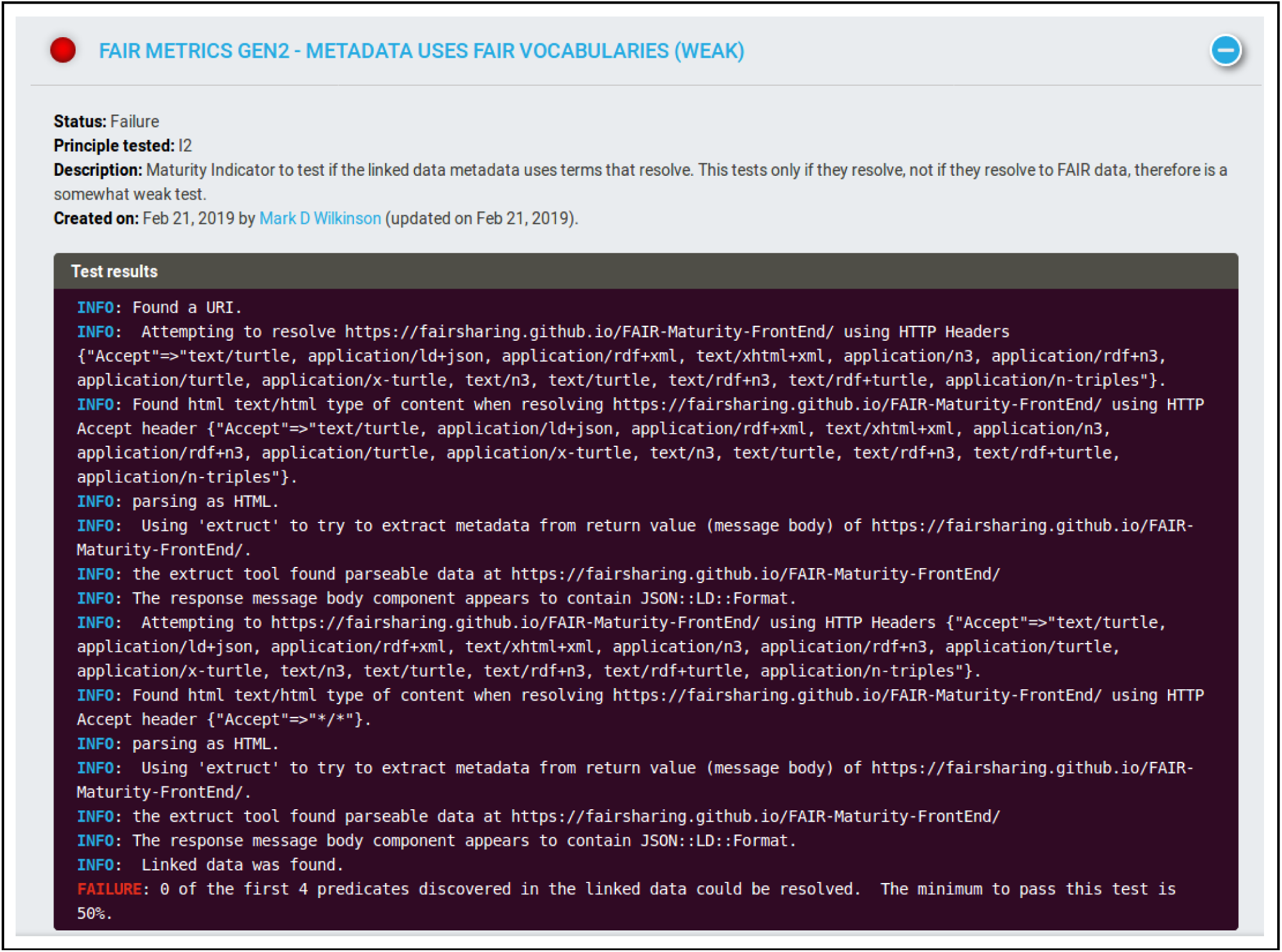
The output of the “Metadata uses FAIR vocabularies (weak)” test. Numerous lines of information indicate the activities that were undertaken by the test. The final line indicates failure of the test, and the reason for that failure.

It is clear, therefore, that we must add metadata to the FAIR Evaluator homepage (for this test, and for other failed tests). There are many ways to accomplish this; however, for this demonstration, we achieved this by embedding a block of JSON-LD into the homepage of the Evaluator. This JSON-LD follows (primarily) a schema.org format, and includes things such as the identifier of the metadata record (by its newly-minted w3id permanent URL), and a pointer to the “data” in the form of a link to the GitHub software repository, using the schema.org *codeRepository* property. The final metadata record is provided in Fig. 4.

**Figure 4.**
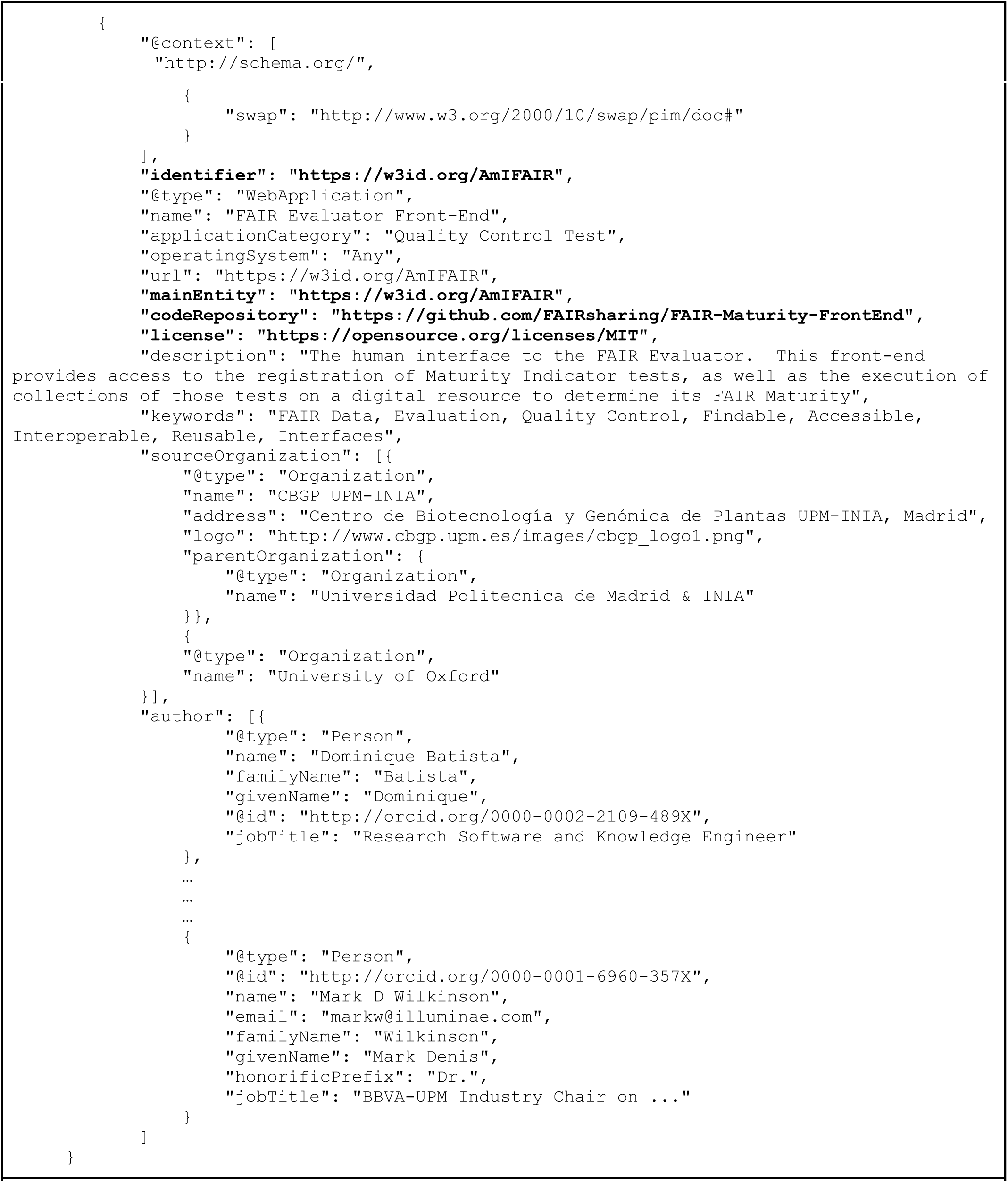
Enriched metadata in the Evaluator page HTML. Important metadata features, required to pass additional MI Compliance Tests, are highlighted in bold. Other metadata features are provided in anticipation of future tests related to “richness” of metadata - i.e., to improve searchability and citability.

Having completed the improvements, we repeated the assessment of the FAIR Evaluator using the same “Maturity Indicator Tests applicable to Software Applications” Collection, and find that now 17 out of the 20 tests pass. There are three tests that our FAIR Evaluator continues to fail:

1. **Data identifier persistence:** Failing this test is expected when the “data” is the software code in the GitHub repository, because GitHub URLs are not a kind of permanent URI. It is possible to create a persistent URI by creating another w3id redirect to the root URL of your software repository.
2. **Searchable in major search engines:** This is an example where the result of a test also relies on a third party tool. Even if we have added the appropriate schema.org discoverability metadata to the FAIR Evaluator homepage, it has not yet been indexed by major search engines. If we run the test in a few days the output will change.
3. **Metadata persistence:** At the moment this test fails because the organization that currently hosts the FAIR Evaluator homepage does not have a persistence policy for Web pages. This is an example of a test that is (usually) outside of the data owner’s control, and they should discuss it with their host.

With only a few improvements, we raised the result of the assessment of our FAIR Evaluator software to its maximal expected score; yet, we still have a clear set of actions that could be applied to improve the score further. However, it is important to note that one should not chase a 100% score at any cost. FAIRness is not a competition, rather, FAIRness refers to a maturation process where digital objects are rendered increasingly self-descriptive to the machine. Communities decide what FAIR practices are most important to them, essentially setting the targets for themselves, allowing members of that community to evolve over time while realistically operating within their budgets in order to achieve their best FAIR performance.

## Methods

The overall workflow for executing FAIR evaluations is summarized in Fig. 5.

**Figure 5.**
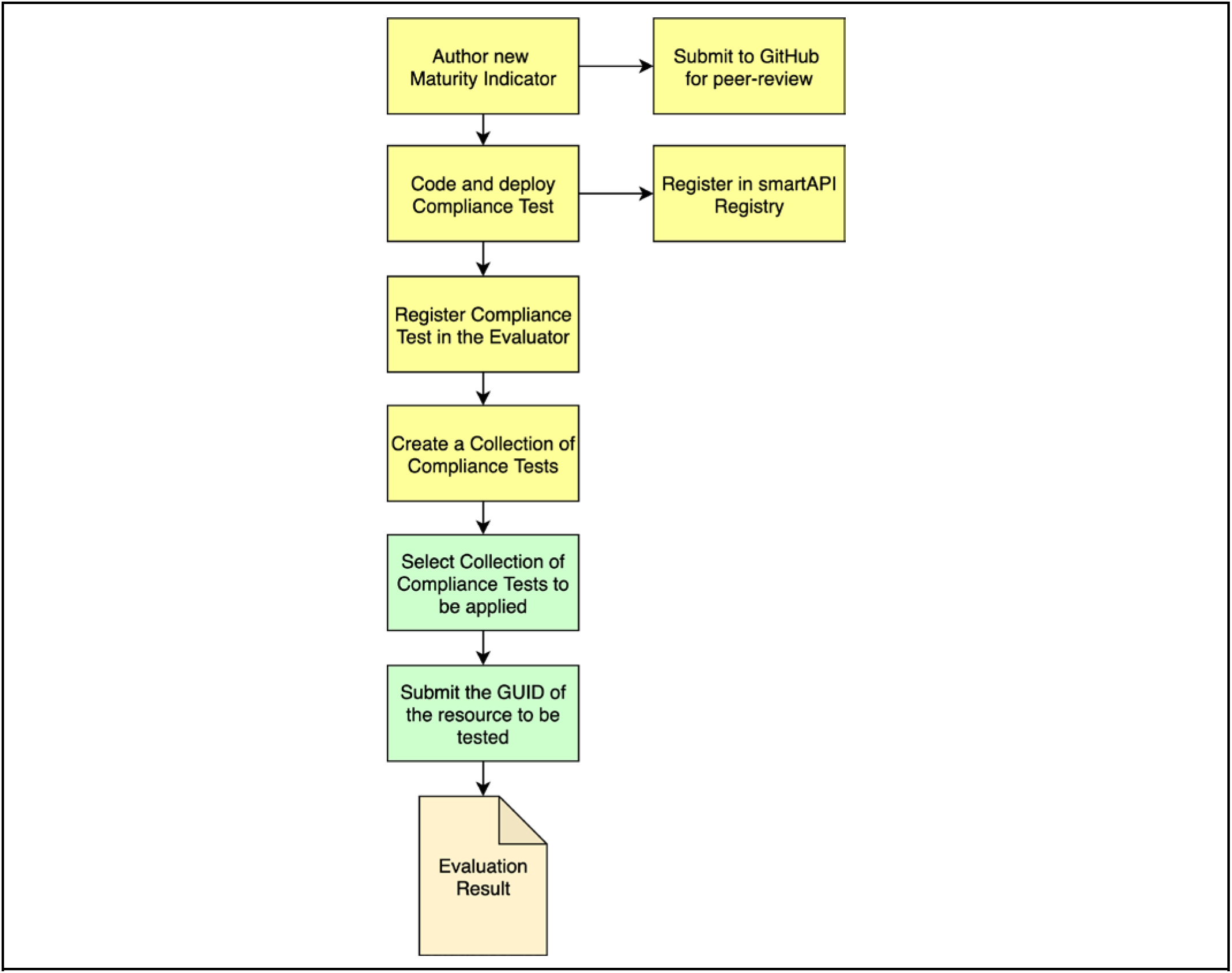
The workflow for executing a FAIR Evaluation. Most steps are optional (yellow boxes), and only become necessary if the set of published MIs and/or Collections is determined to be insufficient. The only steps required for every evaluation are to select a Collection of Compliance Tests appropriate for the digital resource being evaluated, and to submit the GUID of the metadata record for that digital resource (green boxes).

### Creating new MIs

MIs are defined using a prescribed template (https://github.com/FAIRMetrics/Metrics), which captures key aspects such as its unique identifier, the FAIR Principle that it is intended to evaluate, the reason why the evaluated feature is relevant for FAIRness, the precise set of queries/tests that are applied to a resource in order to evaluate its content, and what constitutes success versus failure. A Markdown version of this template is available such that community members may author and submit their own MIs through a pull-request on GitHub. Upon a successful pull, the MI folder is scanned by the standards registry FAIRsharing^8^, and a record of the new MI proposal is then published in a FAIRsharing record, tagged as “In Development”, until finalised when it is marked as “Ready”.

### Nanopublications

In order to make the MIs machine readable, we automatically convert them to nanopublications. Each Markdown-based MI is parsed into its own nanopublication, identified using the *fairmi* namespace (https://w3id.org/fair/maturity_indicator/terms/Gen2/). This includes defining the default human-readable label, linking it to the GUID of the associated FAIR Principle, formally assigning it to the *fairmi:FAIR-Maturity-Indicator* class, and representing the textual descriptions of the MI template as Linked Data predicate-values (e.g. *fairmi:measuring “…”*). Finally, we mint Trusty URIs^9^ to enforce the immutability and verifiability of these nanopublications.

### Creating new compliance tests

A Compliance Test is a Web-based application that tests compliance of a Resource against an MI. Its purpose is to determine if a machine is capable of autonomously discovering the facet or feature of (meta)data that is described by the associated MI. As such, the single piece of information provided to a Compliance Test is a GUID representing a *metadata* record. From that starting information, the test is completed with no additional (manually-provided) data.

Compliance Test interfaces must be described using a smartAPI (Swagger/OpenAPI-based) interface definition^6^, including rich metadata, thus ensuring that the tests themselves are FAIR. It is also recommended, for increased visibility and FAIRness, that these are then also registered in the smartAPI registry (http://smart-api.info/). Further details on the interface of a Compliance Test is provided in the project GitHub repository (https://github.com/FAIRMetrics/Metrics). Compliance Tests are independently published, by any stakeholder, and may be invoked over the Web without any additional infrastructure; however, individual tests, or collections of tests, may also be assembled and invoked using The FAIR Evaluator application.

### The metadata harvester

To provide uniformity to the Compliance Tests, an independent module was designed in Ruby to pursue a wide range of different paths in an attempt to extract metadata about a given GUID. These paths were informed by our initial questionnaire-based study, and the deeper exploration of data provider behaviors that resulted from that. The workflow of the Metadata Harvester is shown in Fig. 6. It currently recognizes four kinds of GUID: DOIs, other Handles, InChI Keys, and URIs. Through a series of content-negotiation and extraction processes, it attempts to find metadata using the metadata provision approaches we identified as in-practice within the data publishing community. It executes the same processes in every case, regardless of the source provider, with the exception of InChI, since the resolution mechanism for these GUIDs is explicitly defined. Both “hash” style metadata (e.g., JSON, Microformat) and Linked Data style metadata (e.g., JSON-LD, RDFa) are collected and stored as separate outputs from the Metadata Harvester. The composite of all harvested metadata, from every step, is passed into each of the Compliance Tests, and used as the material upon which the tests are executed. Note that all data retrieved from the Web is cached for efficiency, and all tests that use the Harvester (i.e., all Gen2 bootstrap tests described in this manuscript) first scan for the record(s) in the cache to reduce network calls. This cache is purged every half-hour to allow users to rapidly re-test changes they might make to their (meta)data.

**Figure 6:**
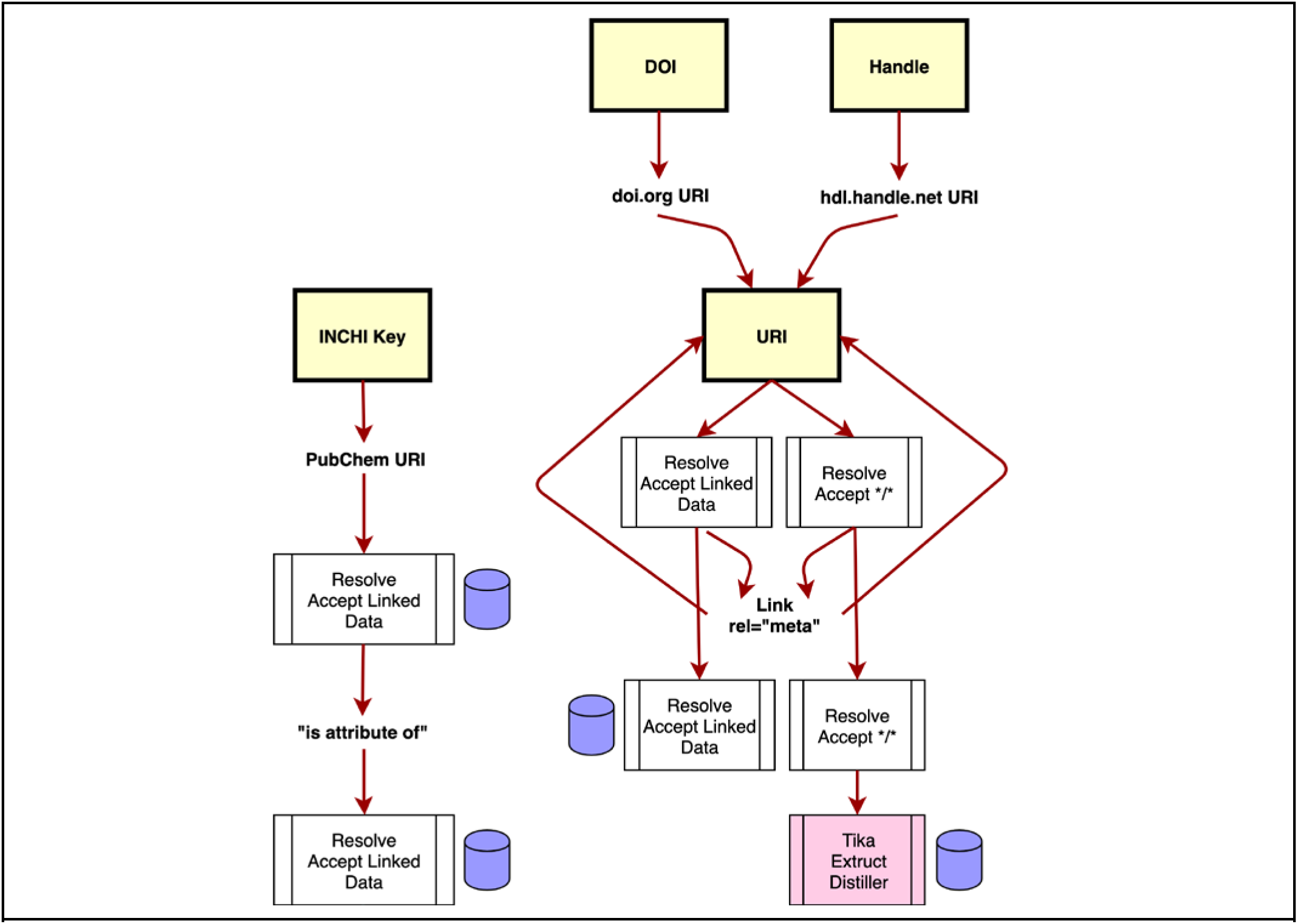
The Metadata Harvester workflow. Yellow boxes are the starting GUIDs, text nodes are some form of URI. Arrows show the flow of information. White boxes are resolution activities, and the Associated Accept headers used. The pink box is a suite of third-party metadata extraction tools. Tika operates on a wide range of non-textual data (e.g. PDFs) to extract embedded metadata. Extruct and Distiller extract embedded metadata within HTML in a variety of formats. Purple barrels are metadata collection steps, where all Linked Data, and hash-style metadata are cached, together with the raw output of the resolution. InChI Keys (left pathway) have a defined two-step resolution mechanism, that supports content negotiation, and are therefore treated as a special case for efficiency. DOIs and other Handles are converted into URIs, and thereafter treated in the same manner. The first step of URI harvesting is to follow all Link rel=“meta” headers, to extract any metadata from these locations following the same workflow as for other URIs. These headers are followed only one layer deep, after which the system returns to the content of the original URI resolution, using content-negotiation.

### The FAIR Evaluator

The FAIR Evaluator is a Ruby on Rails application with four primary components:

- A (read/write/search) registry of known MI Tests and their annotations
- A (read/write/search) registry of collections of MI Tests (“Collections”) and their annotations
- An invocation function, that creates a pipeline of MI Tests and applies them to a resource
- A registry of evaluation results

The FAIR Evaluator is primarily designed for mechanized interaction through its JSON interface; however, a graphical user interface, based on this JSON interface, has been published (https://w3id.org/AmIFAIR) allowing form-based access to all Evaluator functionality. This interface is a client-side Single Page Application written in JavaScript under the AngularJS framework, with source code available through GitHub (https://github.com/FAIRsharing/FAIR-Evaluator-FrontEnd). All figures in the Results section are generated using this interface.

The Evaluator is currently deployed using the Google App Engine (2 CPU, 6G RAM) on an nginx server, with storage being handled by a MySQL 5.7 database on an independent server in the Google Cloud. The Ruby on Rails v5.2.3 configuration is 3 processes with 5 threads each.

### The FAIRsharing service

FAIRsharing is a curated, informative and educational resource on data and metadata standards, inter-related to databases and data policies, with a growing network of users, adopters, collaborators and activities working to enable the FAIR Principles (https://fairsharing.org/communities). Notably FAIRsharing is among the 12 formal flagship outputs of the Research Data Alliance (RDA) (https://www.rd-alliance.org/recommendations-and-outputs/all-recommendations-and-outputs). Its community-roots and relevant content is one of the reasons why FAIRsharing was chosen as the registry of proposed and approved MIs. We are, however, broadening its service in a variety of ways. The core objectives for FAIRness evaluation are objectivity and transparency, as well as automation and scalability. This requires that information against which the results of some of the tests are compared is visible and accessible to all, and it can also be enriched as needed. For example, initially the different available identifier schema were hard-coded into the Compliance Tests, whilst now these are described in FAIRsharing and are retrievable at run-time via its API. New schema and metadata standards will regularly be added to FAIRsharing; its curation team will ensure that new records are described and, if appropriate, inter-linked to databases and repositories that implement them as a way to document their use in the community. Delegating the validation of information about schema and metadata standards to an external service, which can deal with new additions, allows the FAIR Evaluator to adapt to new community standards as they arise without the need to update or re-code the test software. In the future, FAIRsharing will expand to include registration of additional elements required by the Evaluator, such as Web transport protocols, file formats and other standards and policies that are, otherwise, dispersed on the Web.

### Planned work

There are several improvements that could be made to this evaluation platform, and a variety of ways that it could be extended/expanded. For example, the metadata around an evaluation is somewhat thin, and it is difficult to obtain a longitudinal record of the assessment scores for a given resource. A plan is in-place for how to improve this aspect of the Evaluator; however, this will happen only when resources become available to support long-term archival of a potentially very large dataset. In terms of alternative uses for the Evaluator, projects are currently underway to integrate evaluations into data management plans, where tests can be run on existing resources in order to provide advice on what improvements should be included within the plan. The modular nature of this evaluation system, and its ability to be extended by its users, make it well-suited to be used in this manner. Finally, the bootstrap MIs and Compliance Tests described here represent only the most primitive expectations for FAIR digital objects. Passing all of these tests does not indicate that “you are FAIR”! Together with the stakeholder community, we will push to create increasingly challenging tests, in order to incrementally migrate data resources towards more accurate, powerful, and rich automated discovery and reusability behaviors.

## Usage Notes

Code for the FAIR Evaluator and each compliance test is available in GitHub (https://github.com/FAIRMetrics/Metrics/tree/master/MetricsEvaluatorCode). The public interface to the prototype Evaluator software is available for anyone wishing to execute evaluations (https://w3id.org/AmIFAIR). Detailed instructions regarding how to create and submit proposals for new MIs are available in our online documentation (https://github.com/FAIRMetrics/Metrics).

## Discussion

In this data-intensive era, the primary objective of the FAIR Principles is the automated discovery and (re)use of digital objects by machines (even if ultimately for manual analyses). Our proposed evaluation framework focuses on determining the set of FAIR indicators that an automated agent can detect in a digital object. Rather than addressing the question “is this resource FAIR?”, which incorrectly implies that FAIRness is binary^2^, our proposed framework addresses more concrete questions on the features that a digital object provides to facilitate its discovery and reuse by an automated agent. This type of assessment is more objective, task-oriented, reproducible, and scalable than questionnaire-based approaches.

We have initialized the Evaluator system with a starter set of 22 MIs that target generic elements of the FAIR Principles. As such, we believe these MIs will be broadly applicable and likely to be broadly adopted among stakeholders. However, communities are likely to differ in their perception of the value of each of the MIs; thus, this could be reflected in the selective bundling of MIs into community specific Collections. Furthermore we anticipate that, particularly for FAIR Principles focusing on domain-relevant metadata (especially F2 and the R’s), there will be a proliferation of MIs targeting the idiosyncratic domain knowledge and data-reuse needs of individual communities. Stated more directly, not only is the capability to author domain-relevant MIs essential, but for some FAIR facets (such as rich provenance metadata, R1.2) appropriate MIs can only be authored by experienced experts operating in that domain.

Hence, it is essential that the Evaluator system allows for the design of custom-made domain-relevant MIs and their associated Compliance Tests, which complement or complete the starter set we provide. These community-designed elements are also registered, grouped and made discoverable to ensure maximum reuse across-communities, as relevant. This flexibility will help to ensure that digital objects are rendered FAIR under the norms of different communities and at the same time not be ‘unfairly’ tested against requirements that may be considered irrelevant by a that community.

We believe that this is the first general, scalable, automatable framework for FAIR assessments supporting community-defined tests. Its key features are that the tests are transparent, use open standards, and are sufficiently simple to support a broad ecosystem of third-party tooling; the system is modular, and enables a wide range of evaluation approaches, from fully-automated to manual self-assessment; assessments may be rerun at any time in order to support and guide incremental improvements; assessments are reusable objects, and can be run, and examined, by any interested stakeholder, ensuring transparency; the scoring system is a simple binary, with pass/fail scores associated with each test, supporting a wide range of easily-understood reporting and visualization; and most importantly, the feedback provided in the output is designed to positively incentivize and guide the user in what steps can be taken to improve the FAIRness of the digital object.

### Comparison with other evaluation initiatives

A wide range of manually-executed, questionnaire-based evaluation platforms and infrastructures were proliferating contemporaneously with this project, and continue to be developed. A list of these projects has recently been compiled by the RDA FAIR Data Maturity Model Working Group^10^. Some of these assessment frameworks directly refer to a desire to measure specific aspects of FAIRness such as the Research Data Alliance SHAring Reward & Credit (SHARC) IG^11^, The Australian Commonwealth Scientific and Industrial Research Organisation/OzNome 5-star System (http://oznome.csiro.au/5star/), the Dutch Data Archiving and Networked Services ‘FAIR Enough?’ questionnaire (https://docs.google.com/forms/d/e/1FAIpQLSf7t1Z9IOBoj5GgWqik8KnhtH3B819Ch6lD5KuAz7yn0I0Opw/viewform?usp=embed_facebook), and the Australian Research Data Commons FAIR self-assessment tool (https://www.ands-nectar-rds.org.au/fair-tool); others refer to FAIR indirectly through citations, but do not directly ask questions about FAIRness such as the ELIXIR Data Stewardship Wizard (https://app.ds-wizard.org/questionnaire), while others are questionnaires related to data stewardship, without linking the questions to aspects of FAIR such as the Earth Science Information Partners and National Centers for Environmental Information Data Services Maturity Matrix^12^, and the World Meteorological Organization Stewardship Maturity Matrix for Climate Data^13^. This demonstrates the widespread need to provide guidance and tools to the community.

Our approach differs in many ways from above mentioned efforts, primarily because it is strictly focused on detecting and validating behaviors of digital objects that make them machine-readable and reusable, staying true to the FAIR Principles. Other efforts often blur the boundary between FAIRness guidance and data management good practices. For example, some of these initiatives include questions about the curatorial process, repository infrastructure/policy, or stewardship and management plans, therefore measuring something several layers above the digital objects themselves. Although questionnaires serve an important role in improving the overall understanding and appreciation of the research life cycle and the importance of FAIR digital objects, measuring intentions is very distinct from measuring outcomes; and the answers cannot be easily validated. Based on our own experience with Gen 1 and Gen 2 systems, small, unintentional errors may hinder machine-(re)usability, but may go undetected even after self-reported FAIR compliance. Moreover, questionnaires are primarily aimed at someone that deeply knows the data resources, and is therefore capable of answering questions about them - most probably the data steward. For this reason, questionnaires are not suited to being used by third party stakeholders (journals, funding agencies, etc.), who lack the knowledge required to answer such questions. Finally, questionnaires do not scale to the size of today’s datasets and the Web; thus, like with Web indexing, FAIR assessments must rapidly become automatable.

### Call for further community contribution

Individual communities may already have stronger requirements for compliance with the FAIR principles than are reflected in the current starter-set of MIs and Compliance Tests, and we encourage them to reach out to us for assistance in creating their own tests. In particular, however, we felt that the FAIR Principles R1.2 (Detailed provenance) and R1.3 (Compliance with community-specific standards) were outside of the scope of this starting set of MIs. Given that different kinds of digital objects may use different metadata models and provenance (e.g. the provenance of a tomography image from a patient biopsy will be different from the provenance of a pottery artifact in a museum collection), we felt that these should be community-specific. We have, however, put considerable effort to document how new community-driven, bottom-up MIs and associated tests can be designed and incorporated into the FAIR Evaluator. We will continue to foster this sense of community under the umbrella of global FAIR-enabling organizations such as GO FAIR and its Implementation Networks (https://www.go-fair.org/implementation-networks/), RDA groups, as well as other international initiatives and projects.

FAIR is aspirational - a continuum of increasingly rich (meta)data features that support the automated discovery and reuse of important data resources. The availability of a tool that objectively measures critical facets of FAIRness allows both data owners, and third-party stakeholders, to track progress as the global community moves all digital resources towards a world where data and services can be accessed, with increasing precision and efficiency, by the emergent world of computational agents.

## Acknowledgements

MDW is funded by the Isaac Peral/Marie Curie cofund with the Universidad Politécnica de Madrid, Ministerio de Economía y Competitividad grant number TIN2014-55993-RM, and the European Joint Programme on Rare Diseases (H2020-EU 825575). Throughout phase 1 and 2 of the work, S.A.-S., P.MQ., DM and PRS have been funded by grants awarded to S.A.-S. from the UK BBSRC and Research Councils (BB/L024101/1; BB/L005069/1), EU (H2020-EU 634107; H2020-EU 654241; H2020-EU 676559; H2020-EU 824087), IMI (116060; 802750), NIH (U54 AI117925; 1U24AI117966-01; 1OT3OD025459-01; 1OT3OD025467-01, 1OT3OD025462-01), and from the Wellcome Trust (212930/Z/18/Z; 208381/A/17/Z). MD is supported by grants from NWO (400.17.605;628.011.011), NIH (3OT3TR002027-01S1; 1OT3OD025467-01; 1OT3OD025464-01), and ELIXIR, the research infrastructure for life-science data. MP was supported by the UPM Isaac Peral/Marie Curie cofund, and funding from the Dutch Techcenter for Life Sciences DP. LOBS and ES are supported by the Dutch Ministry of Education, Culture and Science (Ministerie van Onderwijs, Cultuur en Wetenschap), Netherlands Organisation for Scientific Research (Nederlandse Organisatie voor Wetenschappelijk Onderzoek), and the Dutch TechCenter for Life Sciences. We thank the NBDC/DBCLS BioHackathon series where many of these MIs and their tests were designed, and we particularly wish to acknowledge the participation of the Dataverse team, especially Julian Gautier and Derek Murphy, at IQSS, Harvard, in addition to Todd Vision from Data Dryad.

## Author Contributions

MDW, MD, SAS, LOBS, and ES designed the Metric and Maturity Indicator rubric and templates, and designed the Gen1 Metrics. MDW, ES, and MC executed the questionnaires and interpreted the results. MDW, MD, SAS, LOBS, ES and MC authored the self-archived report on the Gen1 Metric study. MDW designed the Gen2 MIs, Compliance Tests, and the Evaluator software. MP built access libraries to interact with the Evaluator JSON interface. DB created the Evaluator Front-end. PM and PRS manage the interaction between the FAIR Metrics GitHub and FAIRsharing, and coded new functionality into FAIRsharing to support FAIR evaluations, and tested the framework. TK maintains the GitHub repository of FAIR MIs, and their representation as nanopublications. MDW and SAS wrote the final version of the manuscript. All authors read, edited, and approved the manuscript.

